# H3k27ac-HiChIP in prostate cell lines identifies risk genes for prostate cancer susceptibility

**DOI:** 10.1101/2020.10.23.352351

**Authors:** Claudia Giambartolomei, Ji-Heui Seo, Tommer Schwarz, Malika Kumar Freund, Ruth Dolly Johnson, Sandor Spisak, Sylvan C. Baca, Alexander Gusev, Nicholas Mancuso, Bogdan Pasaniuc, Matthew L. Freedman

## Abstract

Genome-wide association studies (GWAS) have identified more than 140 prostate cancer (PrCa) risk regions which provide potential insights into causal mechanisms. Multiple lines of evidence show that a significant proportion of PrCa risk can be explained by germline causal variants that dysregulate nearby target genes in prostate-relevant tissues thus altering disease risk. The traditional approach to explore this hypothesis has been correlating GWAS variants with steady-state transcript levels, referred to as expression quantitative trait loci (eQTLs). In this work, we assess the utility of chromosome conformation capture (3C) coupled with immunoprecipitation (HiChIP) to identify target genes for PrCa GWAS risk loci. We find that interactome data confirms previously reported PrCa target genes identified through GWAS/eQTL overlap (e.g., *MLPH*). Interestingly, HiChIP identified links between PrCa GWAS variants and genes well-known to play a role in prostate cancer biology (e.g., *AR*) that are not detected by eQTL-based methods. We validate these findings through CRISPR interference (CRISPRi) perturbation of the variant-containing regulatory elements for *NKX3-1* and *AR* in the LNCaP cell line. Our results demonstrate that looping data harbor additional information beyond eQTLs and expand the number of PrCa GWAS loci that can be linked to candidate susceptibility genes.

## Introduction

Prostate cancer (PrCa) is the most common cancer in men, the second leading cause of cancer deaths worldwide, and has a strong familial component ^1–4^. Genome wide association studies (GWAS) have identified single nucleotide polymorphisms (SNPs) associated with increased risk of PrCa across 147 predominantly non-coding genomic loci ^5^, collectively explaining 28.4% of familial risk ^6–9^. With the vast majority of risk variants discovered by PrCa GWAS located outside protein-coding regions, the molecular mechanisms driving pathogenesis have only been described for a handful of loci ^10–15^. GWAS loci predominantly colocalize with tissue-specific regulatory elements thus supporting the hypothesis that risk variants exert their effects on disease by influencing the transcriptional levels of their target genes. For example, a substantial proportion of PrCa heritability lies in regions marked by H3K27ac ^16^, a histone modification marking active enhancers and promoters ^17–19^.

A standard approach to link variants to genes is through expression quantitative trait loci (eQTL) analysis, which identifies genotypes that correlate with transcript levels across individuals in a target tissue of interest ^20–22^. Overlapping eQTLs with variants identified by GWAS is a powerful tool to prioritize target genes for further functional investigation ^20,21,23–28^. However, eQTLs have several limitations. They are primarily studied under steady-state conditions, therefore eQTLs that are elicited under specific conditions are missed. For example, context-specific eQTLs have been identified after stimulation with interferon-γ and bacterial lipopolysaccharides in monocytes ^29^. Second, statistical power for eQTL identification is dependent on sample and effect size with many eQTLs being undetected (i.e. a false negative), particularly for hard-to-collect tissues and/or eQTLs in a rare cell type. Despite some PrCa GWAS loci mapping near well-known prostate cancer biology genes, such as *FOXA1, GATA2, AR*, and *MYC*, the absence of strong eQTL associations with these transcripts is notable. This suggests that traditional eQTL/GWAS overlap may fail to detect important susceptibility genes for PrCa.

Chromosome conformation capture (3C)-based assays have recently emerged as powerful tools to evaluate physical interactions between regulatory elements and target genes in the context of GWAS, including prostate cancer risk ^30–32^ and can be used as an orthogonal method to validate eQTL-based findings. Efficiencies of 3C-based techniques can be increased by enriching for interactions via chromatin immunoprecipitation against proteins, such as H3K27ac, a marker of active promoter/enhancer activity (HiChIP) ^33–35^ or through nucleic acid hybridization^36^.

Notably, across a number of models, enhancer-promoter contacts (loops) pre-exist at stimulation-responsive genes prior to the stimulus ^37–39^. In other words, stimulation does not induce significant *de novo* looping even at genes that are transcriptionally responsive in differentiated cells. These observations raise the hypothesis that looping (measured even in the steady-state) can identify GWAS target genes beyond traditional GWAS/eQTL overlap; that is, looping could reveal eQTL-target gene relationships that are observable only under certain contexts (e.g., cell-type) and/or capture eQTLs with small, but important, effects.

We utilized high-resolution H3K27ac-HiChIP data in the LNCaP prostate cancer cell line to identify links between target genes and PrCa risk loci. We linked 99 out of 130 known susceptibility loci for PrCa with 665 genes and observed a significant overlap between eQTL-target gene pairs and loops. Notably, we identified looping between candidate PrCa causal variants from GWAS and genes with established roles in PrCa biology at eQTL-negative loci. We used CRISPR interference (CRISPRi) to functionally validate the enhancer-promoter interaction for *NKX3-1* and *AR*. Overall, our results confirm that 3C-based strategies can not only validate eQTLs, but can also discover important target gene links not detected by current GWAS/eQTL strategies.

## Results

### A high-resolution H3K27ac-HiChIP chromatin contact map for LNCaP

We performed H3K27ac HiChIP in LNCaP across 5 biological replicates and identified 126,280 loops (FitHiChIP, FDR<0.01, **Methods** and Figure 1). We called between 4,000 and 14,000 loops from a read depth ranging from 183,000 reads to 235,000 reads across individual replicates (**Table S1**). We observed that loop count and loop length depended on read depth. Replicates 1 and 5 had the highest number of high-quality uniquely mapped read pairs and final number of loops, and therefore the highest median loop length and paired end tag (PET) counts (**Figures S1a and S1b**). The intersection of significantly called loops across the 5 replicates ranged from 20% to 60%. Once the individual replicates data was merged, each replicate shared 90% or more significantly called loops with the merged data (**Table S2**). Pairwise correlations of PET counts supporting each loop were significant across all replicates (⍴> 0.7, p-value < 0.001, Figure 2). Merging data across all replicates substantially increased the number of significant loops to 126,280. The mean (median) loop length was 173kb (95kb), and the mean (median) number of paired-end tags (PETs) per loop was 46 (19) (**Table S1**). As expected, the number of loops decreased with increasing loop distance (**Figure S1b**). 17,690 genes (out of 27,063 RefSeq genes considered) had promoters overlapping at last one loop; each gene promoter overlaps a mean (median) of 7.9(6) loops per gene, with a mean (median) of 351 (174) PETs per gene. Gene connectivity, defined as the number of loops per gene promoter (or as total number of PETs per gene), is moderately correlated with gene expression activity (as assayed by RNA-seq in the same cell line, **Methods**) (Spearman ⍴ = 0.489; p-value < 2.2e-16) (**Figures S3a** and **S3b).**

**Figure 1:**
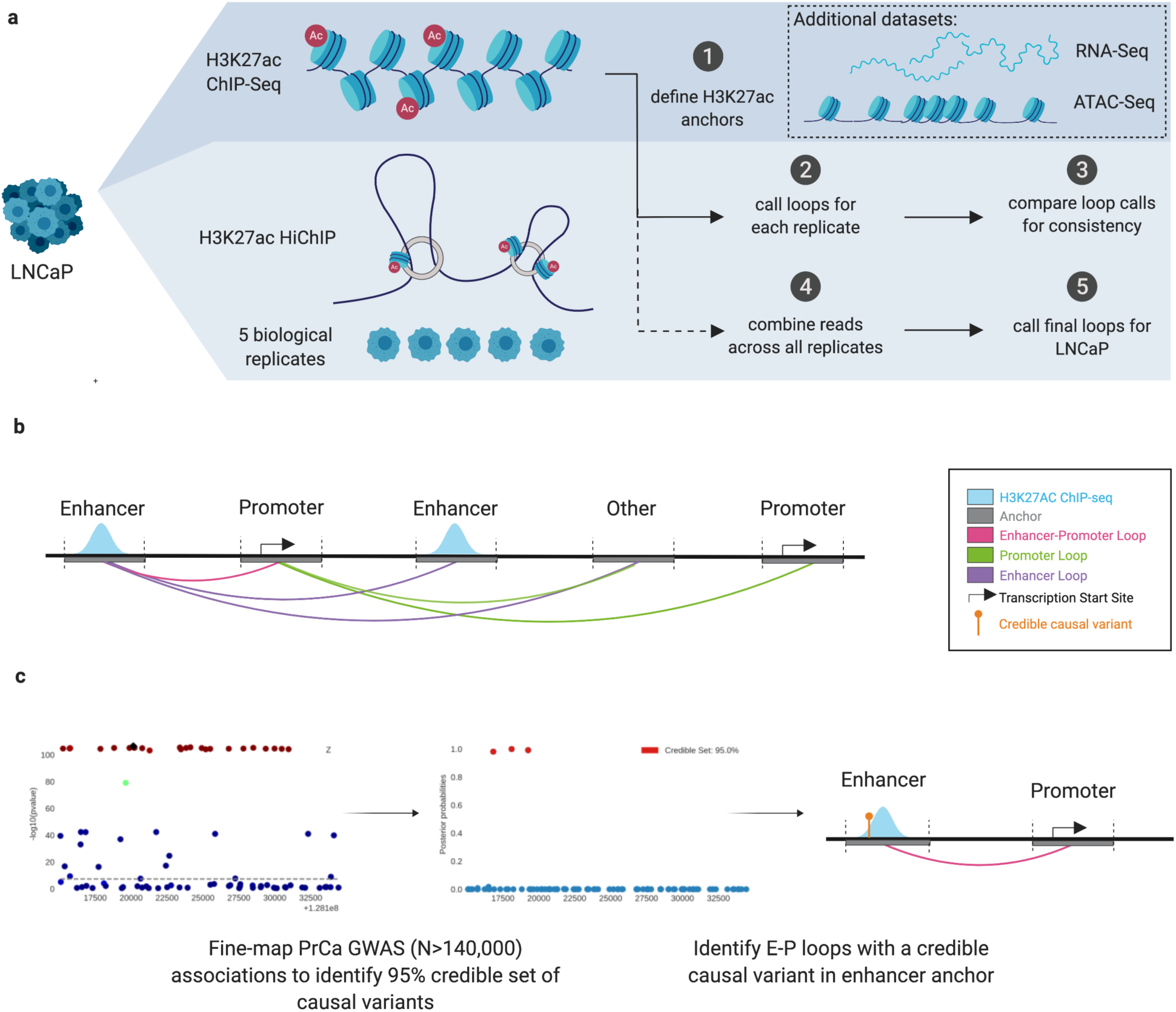
Experimental design. **a.** Description of experimental datasets and workflow of HiChIP analysis to call total LNCaP loops. **b.** Anchor and loop type definitions. **c.** Possible regulatory mechanism describing a causal variant in an enhancer anchor interacting with a promoter of a target gene.

**Figure 2:**
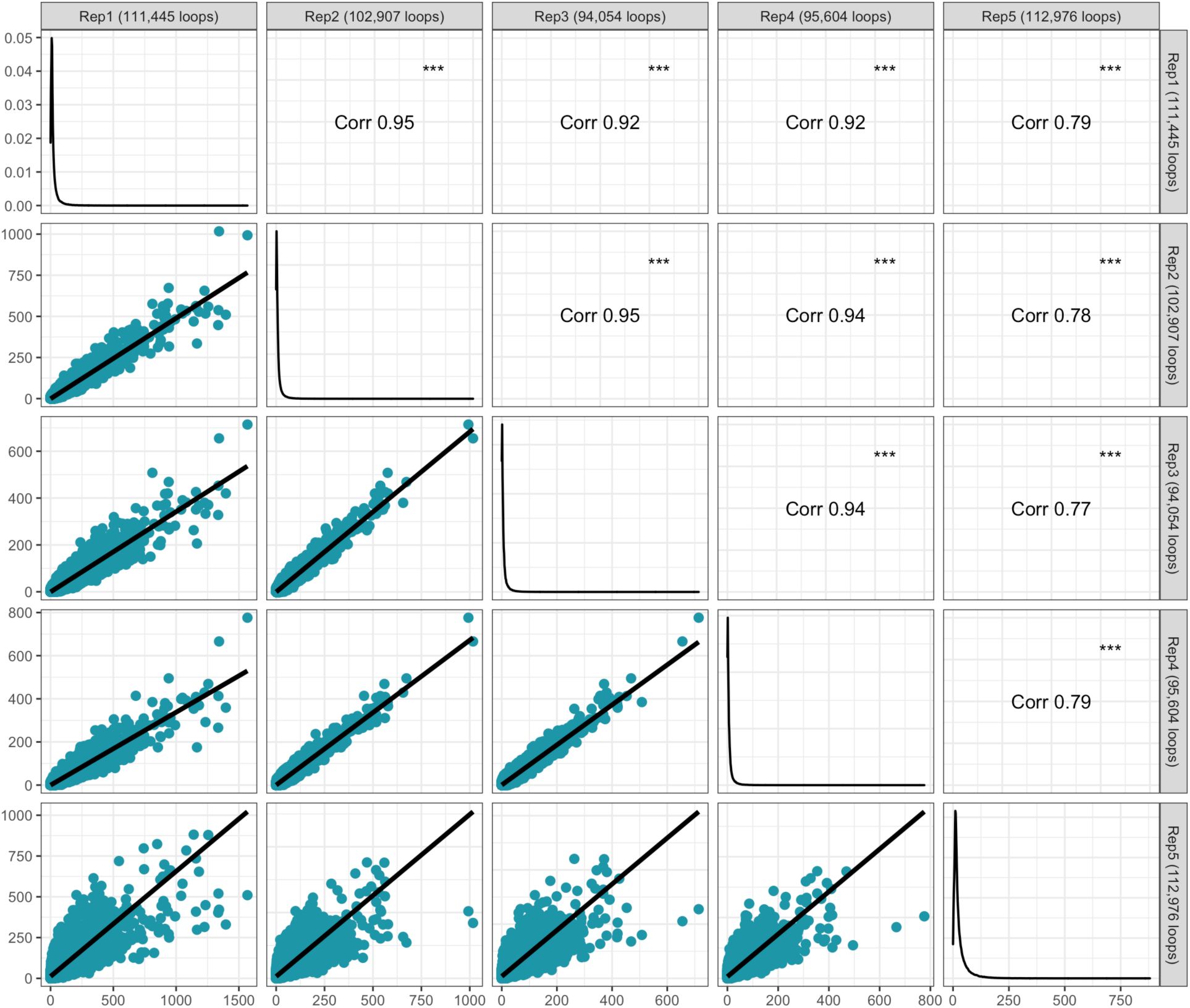
Pairwise correlations of pair-end-tag (PET) counts at loops called significant in at least one of the LNCaP-HiChiP replicates. Every dot represents a HiCHiP loop called at FDR <0.01 in any of the replicates; the union of across replicates N = 114,142 loops. We consider all the loops called at no FDR cut-off. Out of these, we then take the loops called at FDR 0.01 in any of the replicates (i.e. the union of loops across replicates, N = 114,142 loops). Within the bottom left panels, the axis represent the number of pair-end-tags observed in the two compared replicates. The diagonal panel shows the distribution of each replicate’s PET counts. Pearson correlation coefficients are shown on the top right panels. If the PET count is missing for a loop in one replicate but it is present in the other, it is treated as 0 in the correlation estimation. See Supplementary Figure 1 for a comprehensive comparison across replicates.

Prior studies have demonstrated that looping interactions do not qualitatively change in response to a perturbation ^37–39^. To assess whether loop formation correlates with differential expression across perturbations and states, we evaluated the looping status of genes in androgen stimulated cells and during tumorigenesis (**Methods**). We observed that genes that are destined to change under these conditions have pre-existing loops (e.g., upon androgen stimulation, genes with >1.5-fold changes in transcript levels after 4h have 10 loops on average whereas the average number of loops of any gene in LNCaP is 9, see **Table S3; Table S4**). This observation suggests that *de novo* loop formation is not a major mechanism underlying gene expression changes in this data.

### H3K27ac-HiChIP loops are enriched for eQTLs in prostate tissue

To evaluate if chromatin interactions are enriched in prostate eQTLs, we assessed the overlap of loops with eQTLs measured in two prostate datasets: Thibodeau eQTLs in prostate normal tissue (n=471) ^40^, and TCGA eQTLs in prostate tumor tissue (n=378) ^41^. Local eQTLs (within 3Mb of every gene) were called using standard approaches (**Methods**). First, we found that the top eQTL variant for each eGene is 2 times more likely to fall within a HiChIP anchor than a random SNP in a 3Mb region centered on the eGene (empirical p-value 0.0099, **FigureS4a**). Second, we found that loops with eQTLs in one anchor were more likely to loop to the promoter of their eGene than expected by chance (1.2-fold change compared to random, empirical p-value = 0.0099, **Methods**), across a range of distances from their target promoters (**Figures S4b** and **S4c)**.

### H3k27ac-HiChIP loops link prostate cancer risk variants to target genes

Next, we quantified the enrichment of GWAS signals at the anchor regions of the loops using stratified linkage disequilibrium score (LDSC) regression analysis ^42^ integrating the H3K27ac-HiChIP loops with the largest PrCa GWAS to date ^5^. Variants residing in loop anchors show high enrichment of prostate cancer heritability as compared to random variants in the genome overall (s-LDSC enrichment of 2.61, p=1.06E-09), and the enrichment is even greater when the variants are also falling within H3K27ac peaks (our calling allows for one anchor not to be acetylated thus yielding many loops with one non-acetylated anchor, see **Methods**) (s-LDSC enrichment of 14.06, p = 2.54e-5 for all loop types, **Table S5**). This confirms that H3K27ac-HiChIP looping localizes relevant PrCa GWAS signal. Interestingly, we observed a higher enrichment when restricting loops to specific categories (e.g., 18.72 for enhancer-promoter, p=8.93e-6, **Table S5**).

Next, we performed probabilistic fine-mapping of the prostate cancer GWAS at 130 of the 147 previously reported risk loci except 8q24 for which exhaustive fine-mapping is available ^11^. We integrated 1,953 HiChIP loops with at least one anchor overlapping a PrCa credible causal variant in 104 risk loci regions. Overall, 2,016 PrCa credible causal variants linked to 665 genes across these 104 loci (Figure 3, **Table S6**). 48 (out of 104) regions have 3 or fewer HiChiP genes overlapping credible GWAS SNPs (**Table S7**). Of the 665 genes, 37 are eGenes in prostate tissue and have evidence of GWAS/eQTL overlap with PrCa risk variants either through TWAS and/or co-localization (Figure 3, Table 1, **Table S8**). One example is *MLPH* (Figure 4a), which is an eGene in prostate tissue and has been previously reported in normal prostate tissue from GTEx ^43^. By contrast, 4 genes (*CEACAM21, MOB2, ASCL2, GDF7*) are eGenes and show evidence of TWAS/Colocalization but are not supported by any HiChiP loop (**Table S8**).

**Table 1:**
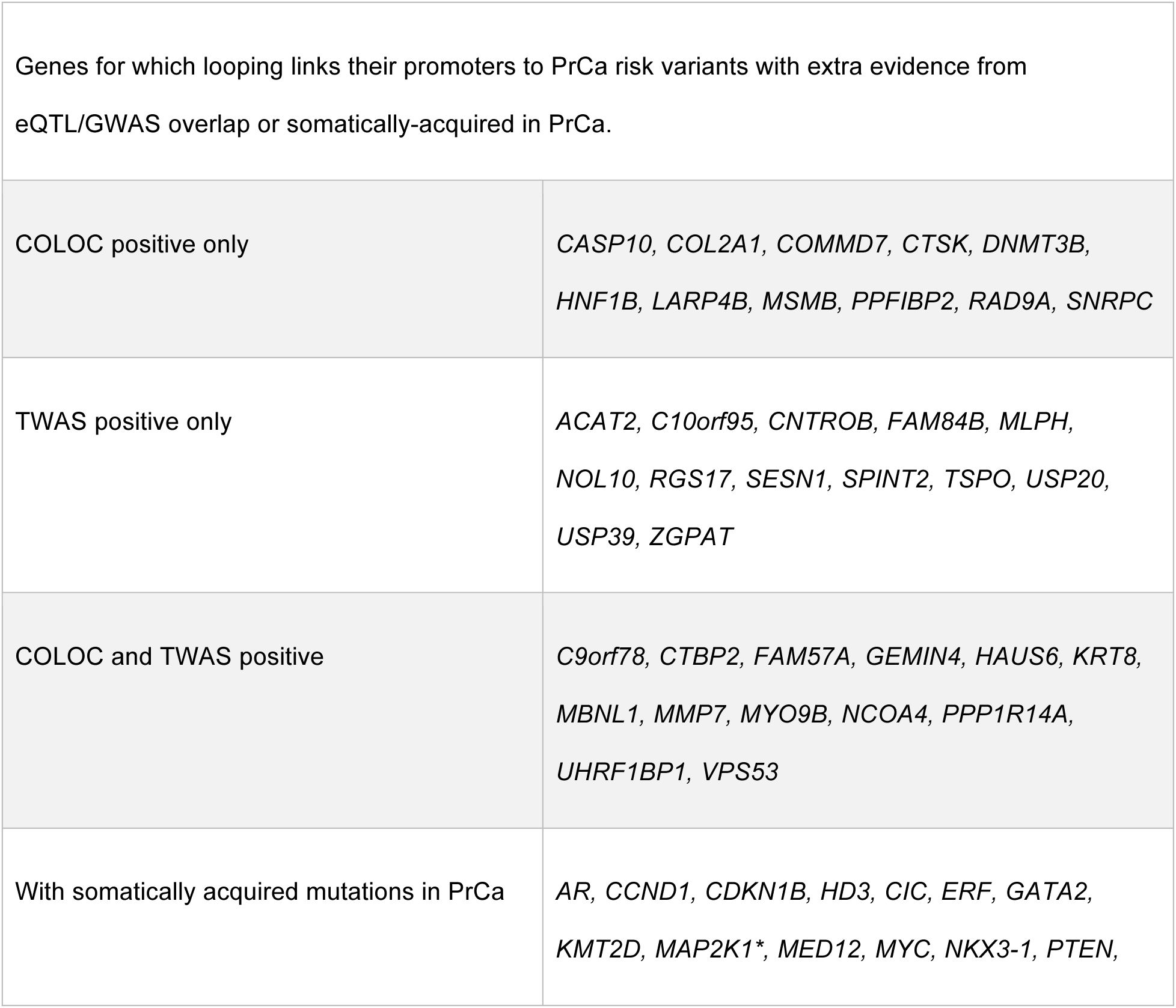
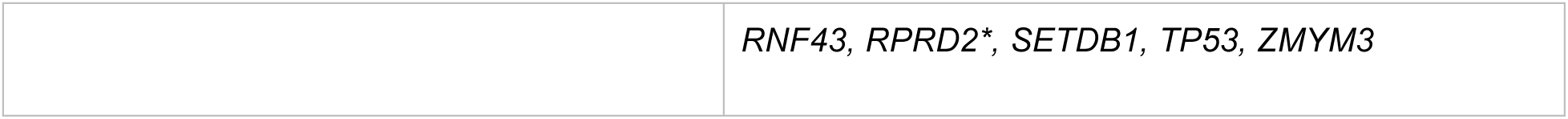
Genes where HiChIP looping data links their promoters to PrCa risk variants. Out of the 665 genes linked by looping to PrCa risk variants (Figure 3), we list genes with evidence of eQTL/GWAS overlap (37) or somatically-acquired mutations in PrCa (18); see **Table S6** for all RefSeq genes. For 37 genes looping links their promoter to PrCa GWAS variants, and also show evidence of eQTL/GWAS overlap through colocalization (COLOC) and/or transcriptome-wide association (TWAS). 18 out of the 119 genes previously reported to have somatically acquired mutations in PrCa show loops linking their promoters to PrCa germline risk variants. The asterisk points to eGenes.

|  |  |
| --- | --- |
| Genes for which looping links their promoters to PrCa risk variants with extra evidence from eQTL/GWAS overlap or somatically-acquired in PrCa. |  |
| COLOC positive only | *CASP10, COL2A1, COMMD7, CTSK, DNMT3B, HNF1B, LARP4B, MSMB, PPFIBP2, RAD9A, SNRPC* |
| TWAS positive only | *ACAT2, C10orf95, CNTROB, FAM84B, MLPH, NOL10, RGS17, SESN1, SPINT2, TSPO, USP20, USP39, ZGPAT* |
| COLOC and TWAS positive | *C9orf78, CTBP2, FAM57A, GEMIN4, HAUS6, KRT8, MBNL1, MMP7, MYO9B, NCOA4, PPP1R14A, UHRF1BP1, VPS53* |
| With somatically acquired mutations in PrCa | *AR, CCND1, CDKN1B, HD3, CIC, ERF, GATA2, KMT2D, MAP2K1\*, MED12, MYC, NKX3-1, PTEN,* |
|  | *RNF43, RPRD2\*, SETDB1, TP53, ZMYM3* |

**Figure 3:**
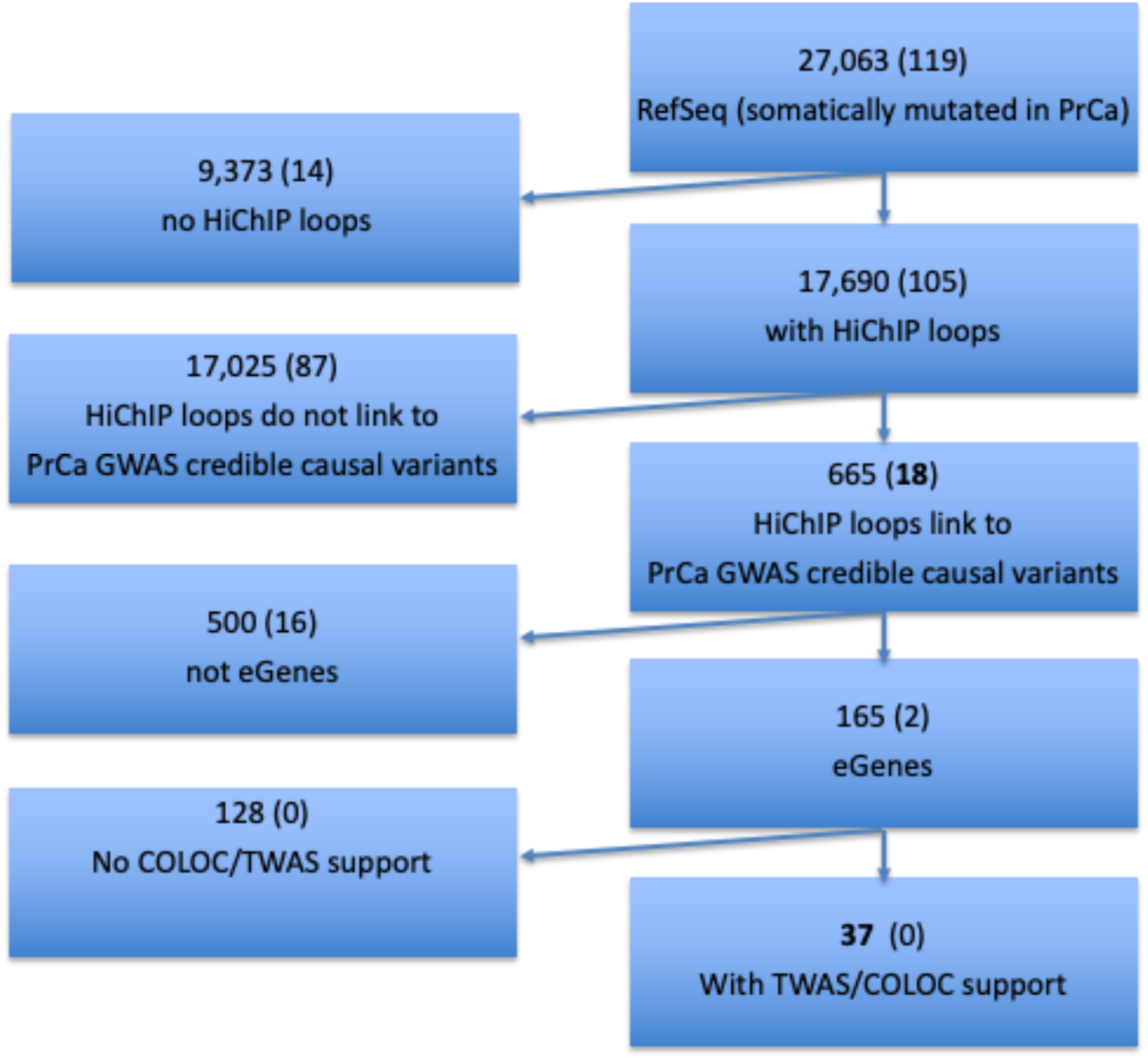
Breakdown of genes with various forms of evidence for linked to PrCa risk variants. Out of 27,063 genes in RefSeq, 17,690 show a HiChIP loop overlaying their promoter. 665 genes at 104 PrCa GWAS regions have a loop linking their promoter to a PrCa credible causal variant. 165 (out of 665) are also eGenes in tissues relevant to PrCa with 37 also showing evidence of co-localization and/or transcriptome-wide association. The numbers in parentheses showcases the breakdown of the 119 genes with evidence of somatically acquired mutations in prostate cancer (**Methods**).

**Figure 4.**
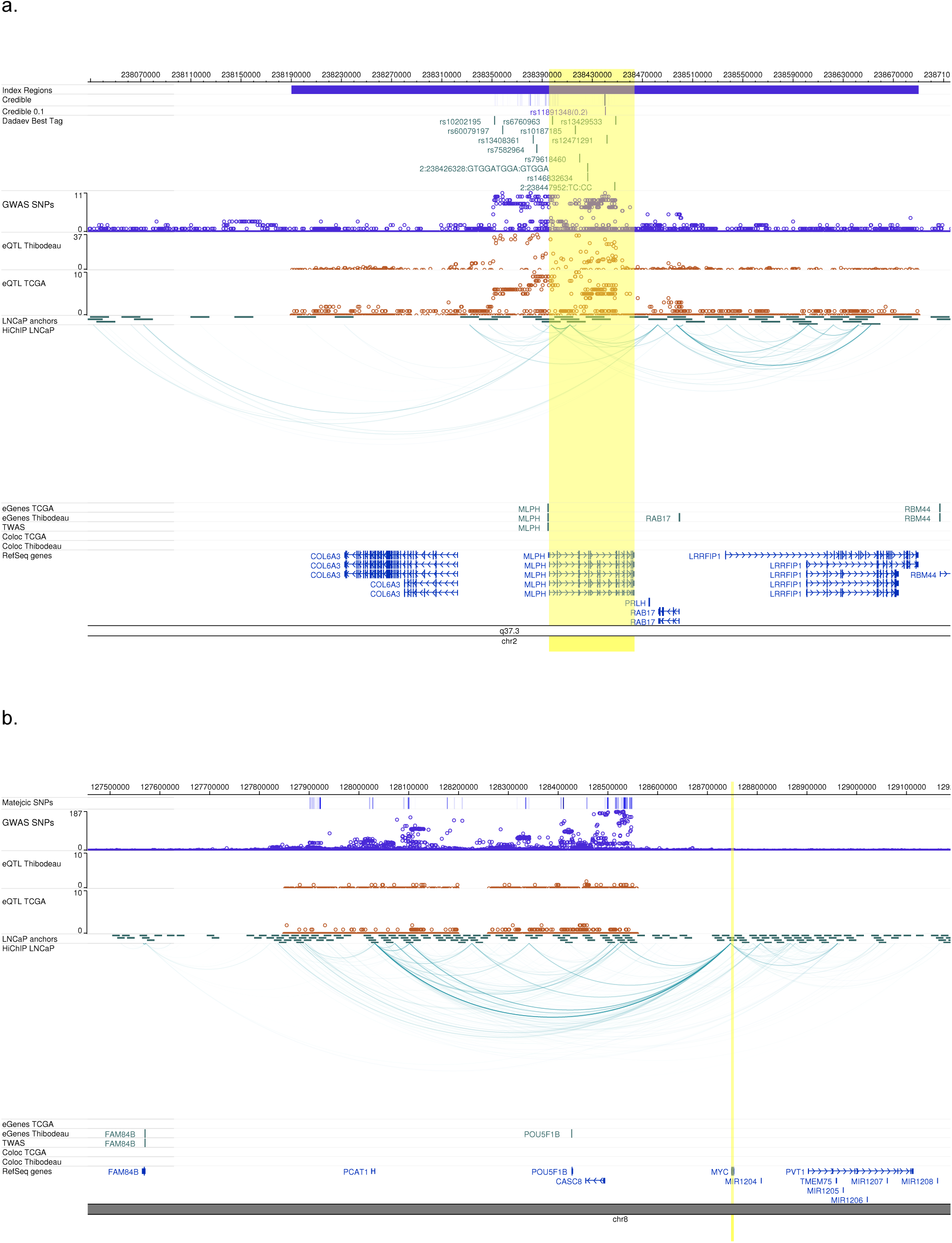
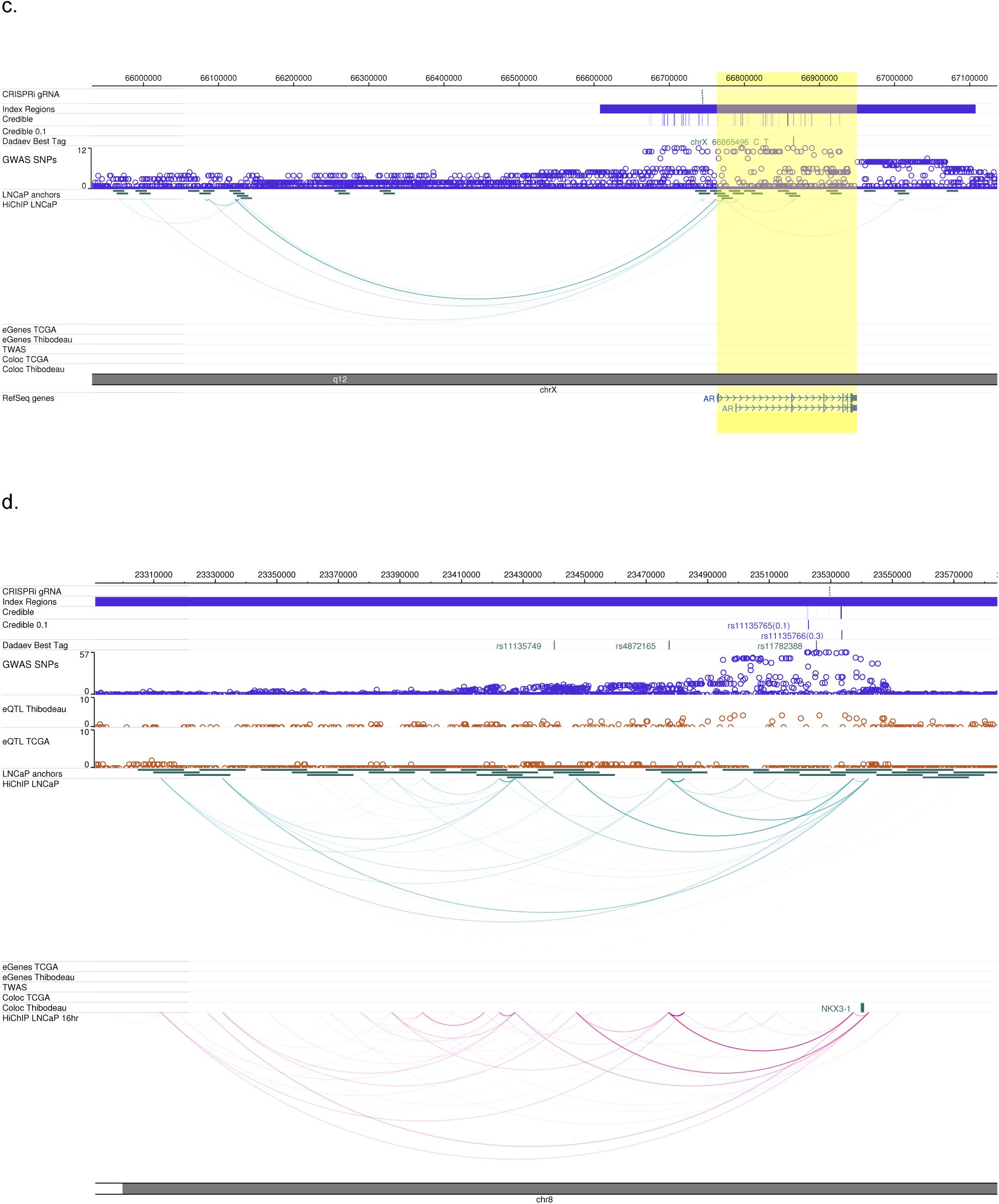
Examples of genes supported by HiChIP loops and GWAS: **a.** *MLPH*, with evidence of eQTLs in prostate tissue using both prostate tissue datasets (Thibodeau and TCGA); **b.** *MYC* and **c.** *AR*, with no evidence of eQTLs. **d.** *NKX3-1*, with weak evidence of eQTLs in prostate tissue. The tracks shown are listed below. <u>**Index Regions**</u>. the fine-mapped region. <u>**Credible.**</u> Position of the SNPs included in the 95% credible SNP set of being the most likely causal variants from PAINTOR analyses (Methods). The color of the track is deeper for SNPs with higher probability of being causal from 0 to 1. <u>**Matejcic SNPs**</u>. Position of fine-mapped SNPs from Matejcic et al. ^11^ <u>**CRISPRi gRNA**</u>. gRNA positions targeted for the CRISPRi experiment. <u>**Credible 0.1.**</u> Position and name of SNPs that reach a posterior probability of being causal of 0.1. <u>**Dadaev Best Tag**</u>. Position and name of SNPs within 341 "best tag" SNPs fine-mapped in ref. Dadaev et al. <u>**GWAS SNPs.**</u> Each hollow circle represents a SNP; y-axis is the position and x-axis is the −log10(P) genome-wide association with PrCa risk. <u>**LNCaP anchors**</u>. Regions of the genome containing HiChIP anchors. <u>**HiChIP LNCaP.**</u> Loops from merged data of 5 replicates. <u>**eQTL TCGA**</u>. Same as above track. Associations were run on gene expression and genotype data on 494 prostate cancer patients (ref. Pancan website). <u>**eGenes Thibodeau.**</u> Highlighted genes that are eGenes in Thibodeau dataset. <u>**eGenes TCGA**</u> Highlighted genes that are eGenes in TCGA dataset. <u>**eQTL Thibodeau**</u>. Each hollow circle represents a SNP; y-axis is the position and x-axis is the associations between SNP and gene expression (-log10(pvalue)) for SNPs within 3Mb window around a gene promoter. Associations were run using gene expression and genotype data from 471 samples from normal prostate tissue (ref. Thibodeau). <u>**eQTL TCGA**</u>. Each hollow circle represents a SNP; y-axis is the position and x-axis is the associations between SNP and gene expression (-log10(pvalue)) for SNPs within 3Mb window around a gene promoter. Associations were run using gene expression and genotype data from 471 samples from normal prostate tissue (ref. Thibodeau). <u>**TWAS**</u>. Highlighted genes that have a significant TWAS signal in Prostate cancer. <u>**Coloc Thibodeau**</u>. Highlighted genes that have a significant COLOC signal in Prostate cancer, using Thibodeau eQTL data. <u>**Coloc TCGA**</u>. Highlighted genes that have a significant COLOC signal in Prostate cancer, using TCGA eQTL data. <u>**RefSeq genes.**</u> Known human protein-coding and non-protein-coding genes taken from th 2019 RefSeq Release. <u>**HiChIP LNCaP 16hr.**</u> HiChIP loops measured after 16 hours from androgen stimulation.

### Looping maps PrCa GWAS variants to known PrCa biology genes

We noted that many genes previously implicated in PrCa biology, such as *GATA2, AR, MYC*, and *NKX3-1 are nearby* PrCa GWAS risk loci. Despite the clear role of these genes in PrCa, compelling evidence linking risk variants to these genes through expression-based methods is currently lacking. Interestingly, looping provides links from promoters of these genes to GWAS risk regions. We highlight three such genes *MYC*, *AR* and *NKX3-1*.

*MYC* has an uncontested role in cancer biology and has been associated with numerous cancer types through GWAS ^44–46^, *MYC* has never been reported as an eGene. A study from Matejcic et al. ^11^ fine-mapped the prostate cancer susceptibility region at 8q24 where *MYC* is located and observed 174 variants in the 95% credible set. 169 of these candidate causal variants overlap one of 225 HiChIP loops, and the majority (152/169) link to the *MYC* promoter (Figure 4b, **Table S9**).

*AR* is critical for prostate cancer as PrCa is dependent on the actions of androgens and therefore on the function of the *AR* gene. More than half of primary tumors and almost all tumor metastases are associated with overexpressed or deregulated *AR* ^47,48^, and mutations in the *AR* gene have been associated particularly with tumor progression ^49–51^. We identified H3K27Ac HiChIP loops that link PrCa credible SNPs located in the X chromosome with the *AR* promoter, potentially pointing to enhancer regions important for PrCa (Figure 4c). Moreover, these variants have never been implicated as eQTLs.

*NKX3-1* is another interesting example since it is one of the most androgen responsive genes in the LNCaP cell line, it is involved in prostate development, and is recurrently mutated in advanced PrCa ^52–55^. We measured looping before and after androgen stimulation in the LNCaP cell line and we observed that the number of *NKX3-1* promoter loops is largely unchanged (**Methods, Table S4** and Figure 4d). For example, the expression changes by ~2 fold after 4(16) hours of androgen stimulation (logFC = 2.23(1.88) p-value < 1E-50), while the total number of loops change minimally (from 14 loops to 11 and to 13, respectively for the two time points, **Table S4**). These data indicate that transcriptionally dynamic genes, which may represent context-dependent eQTL targets are discoverable through looping.

### Looping identifies germline-somatic interactions

We next investigated germline-somatic interactions by evaluating whether genes known to be somatically mutated in PrCa oncogenesis also show evidence of looping to germline PrCa GWAS. We identified a set of 119 prostate cancer–genes curated from large-scale PrCa studies that show evidence of somatically acquired mutations ^56–58^ (**Methods**). Interestingly these genes are on average closer to a PrCa credible causal variant when compared to other genes within a 3Mb region, providing additional evidence of their importance to PrCa (**Figure S5**, 30% of PrCa genes compared to 8% of all genes are within 100Kb of PrCa credible causal variant). Eighteen (out of 104) genes have HiChIP loops linking a credible causal variant to the promoter of the gene (Table 1). Strikingly, none show significant colocalization GWAS/eQTL, with only 2 genes being eGenes in existing prostate transcriptomic data (Figure 3).

### Validation of enhancer-promoter contacts using CRISPRi

Next, we validated candidate HiChIP SNP-gene targets at PrCa two GWAS loci using CRISPRi in LNCaP cells. We selected two GWAS loci where anchors containing a candidate causal variant looped to genes that play clear roles in PrCa biology, *NKX3-1* and *AR* (Figure 5). As described above, these two loci have not been strongly implicated through eQTL- or COLOC/TWAS-based analyses. Putative regulatory enhancer regions containing candidate causal variants within a HiChIP anchor and overlapping DNase hypersensitivity sites (DHS) were identified and targeted. Guide RNAs (gRNAs) were designed and tested against distinct DHS peaks to evaluate the role of these enhancers on *NKX3-1* and *AR* expression (**Table S10)**. Suppression of these enhancer regions containing candidate causal variants significantly reduced RNA levels in the target genes (Figure 5, **Table S11**).

**Figure 5.**
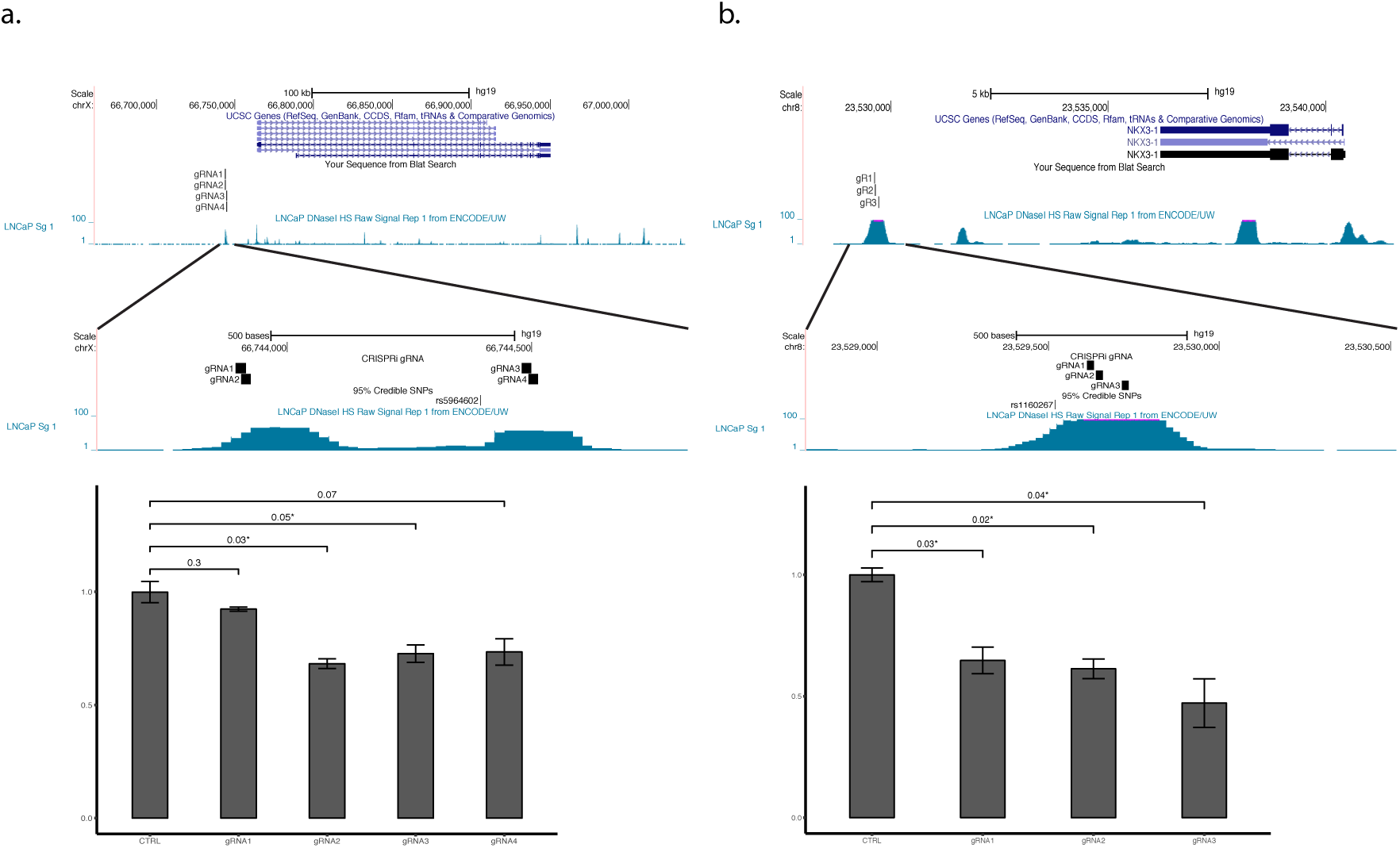
CRISPRi functional validation of the AR and NKX3-1 loci. Functionally relevant enhancers were identified by integrating genetic (PrCa causal variants) and epigenetic datasets (DHS peaks and HiChIP anchors in LNCaP cell line). **a.** AR gene genomic region, LNCaP DHS signals, 95% credible SNP set and gRNA positions targeting DHS peaks for the CRISPRi experiment. Bottom panel shows the suppression effect on the AR gene using four different gRNAs compared to the non-human targeting negative control. Three out of the four tested gRNAs showed ~1 fold significant suppression on the AR gene expression. Columns demonstrate averages of two biological replicates, error bars represent standard errors. **b.** NKX3-1 genomic region, LNCaP DHS signals, 95% credible SNP set and gRNA positions targeting DHS peaks for the CRISPRi experiment. Bottom panel shows the suppression effect on the NKX3-1 gene using three different gRNAs compared to the non-human targeting negative control. All three tested gRNAs showed ~1 fold significant suppression on the NKX3-1 gene expression. Columns demonstrate averages of two biological replicates, error bars represent standard errors.

## Discussion

A central issue driving post-GWAS studies is a mechanistic understanding of non-protein coding risk loci, which account for over 90% of GWAS variants. In this work, we outlined a systematic approach, based on chromosome conformation capture technology, to link GWAS PrCa risk SNPs to candidate target genes. We used H3K27Ac HiChIP methodology as a means to assay genome-wide enhancer-promoter chromatin interactions. 3C-based methods measure physical interactions and, thus complement other approaches, such as eQTLs, which are based on association between genotypes and transcript levels. eQTL studies can be confounded by LD and are dependent on sample size whereby interactome maps do not suffer from these limitations.

An additional limitation of large-scale expression-based studies is that they are based on steady-state transcript levels. By contrast, studies have demonstrated that looping is less dynamic in response to defined perturbations ^37,39,59^. Stated another way, looping identifies the *potential* of an enhancer-promoter interaction to be active and is less informative as a quantitative readout of transcriptional levels of genes. This observation raises the provocative notion that looping can identify latent, stimulus- and context-dependent eQTLs and highlight important candidate genes without requiring experiments that directly measure these other conditions. This rationale is consistent with recent reports showing that steady-state eQTLs are insufficient to explain the majority of disease heritability ^60^. Indeed, our results showed that important PrCa biology genes interact with risk loci that previously escaped detection through expression-based methods.

Other techniques can be used to assay enhancer-promoter interactions. Traditional Hi-C can assay all loops; however Hi-C can lack resolution in mapping the enhancer-promoter link if not sequenced to extremely high depths. Through enrichment based on immunoprecipitation of a target protein, H3K27Ac HiChIP is an efficient way to detect enhancer-promoter loops and has been shown to identify a similar number of loops with ten-fold less sequencing compared to Hi-C ^61^. ChIA-PET is another whole genome 3C-based assay able to detect protein-centric long range contacts, however this method still requires hundreds of millions of cells per experiment and is less efficient than HiChIP ^61^.

As with any method, there are limitations of HiChIP. First, all 3C-based methods have limited power to confidently detect nearby genes. Second, the computational pipelines to analyze HiCHiP data are still in their infancy and future developments could affect results. HiChIP loop-calling is dependent on H3K27ac ChIP-seq peak-calling used for loop anchors. In this study, we defined promoters based on the longest transcript from TSS from RefSeq, which reports the most representative initiation site across different cell types. Using this definition, novel genes with as-yet unannotated start sites are missed. Furthermore, a 5-kb resolution of the HiChIP data analysis limits our definition of anchors, and there is the risk that the additional 5-kb padding added to anchors could result in decreased resolution of enhancer-promoter links. In this work we focused on H3K27ac histone modification. In future studies, assays using other modifications (e.g., H3K4me1, H3K4me2) could be used in combination to better refine enhancer and promoter regions ^62^.

We demonstrate the benefit of using HiChIP to both validate eQTLs as well as prioritize genes for PrCa that are missed by eQTL-based methodologies. Moving forward, we propose to utilize the complementary techniques of eQTL-based methodologies and HiChIP to prioritize genes at GWAS loci. This work also creates an opportunity to create a unified model combining expression and interaction information to extract the strengths of both methods.

## Methods

### H3K27ac ChiP-Seq in LNCaP

H3K27Ac ChiP using either LNCaP (CRL1740, ATCC) was performed as previously described ^51^. 10 million cells were fixed with 1% formaldehyde at room temperature for 10 minutes and quenched. Cells were collected in lysis buffer (1% NP-40, 0.5% sodium deoxycholate, 0.1% SDS and protease inhibitor (#11873580001, Roche) in PBS) ^63^. Chromatin was sonicated to 300-800 bp using Covaris E220 sonicator (140PIP, 5% duty cycle, 200 cycle burst). H3K27ac Antibody (C15410196, Diagenode, 1:600 ratio) were incubated with 40 µl of Dynabeads protein A/G (Invitrogen) for at least 6 hours before immunoprecipitation with the sonicated chromatin overnight. Chromatin was washed with LiCl wash buffer (100 mM Tris pH 7.5, 500 mM LiCl, 1% NP-40, 1% sodium deoxycholate) 6 times for 10 minutes. Eluted sample DNA wes prepared as the sequencing libraries using the ThruPLEX-FD Prep Kit (Rubicon Genomics). Libraries were sequenced using 150-base pair single reads on the Illumina platform (Illumina) at Novogene. For further analyses, we used the union of ChipSeq narrow and broad peaks in regular media. This comprises 49,638 ChipSeq peaks in LNCaP cells (length of peaks ranged from 146 to 129,126 bases). 43,335 out of 49,638 peaks overlap HiChIP anchors.

### Peak calling

ChIP-seq and ATAC-seq were processed through the ChiLin pipeline ^64^. Briefly, Illumina Casava1.7 software used for base calling and raw sequence quality and GC content were checked using FastQC (0.10.1). The Burrows–Wheeler Aligner (BWA, version 0.7.10) was used to align the reads to human genome hg19. Then, MACS2 (v. 2.1.0.20140616) was used for peak calling with a FDR q-value threshold of 0.01. Bed files and Bigwig files were generated using bedGraphToBigWig for H3K27Ac. The union of H3K27Ac narrow and broad peaks was used in the downstream analyses. The following quality metrics were assessed for each sample: 1. Percentage of uniquely mapped reads, 2. PCR bottleneck coefficient to identify potential over amplification by PCR, 3. FRiP (fraction of non-mitochondrial reads in peak regions), 4. Peak Number, 5. Number of peaks with 10-fold and 20-fold enrichment over background, 6. Fragment size, 7. Percentage of the merged peaks with promoter, enhancer, intron, or intergenic, and 8. Peak overlap with DNaseI hypersensitivity sites. For datasets with replicates, ChiLin calculates the replicate consistency with two metrics: 1. Pearson correlation of ChIP-seq reads across the genome using UCSC software wigCorrelate after normalizing signal to reads per million and 2. percentage of overlapping peaks in the ChIP replicates.

### RNA-Seq in LNCaP

1 million of LNCaP cells were harvested for RNA seq. For the DHT treatment samples, LNCaP cells were grown in hormone depleted media for 3 days then treated with 10nM DHT for either 4h or 16h. Total RNA was collected from the cells using RNeasy kit (74104, Qiagen) with DNase I treatment (79254, Qiagen). The library preparations, quality control and sequencing on a HiSeq platforms with paired-end 150 bp (PE 150) were performed by Novogene. RNA-seq data were processed using the VIPER pipeline ^65^. Reads were aligned to the hg19 human genome build with STAR ^66^. FPKM values were calculated with Cufflinks ^67^ for 20,114 RefSEQ genes included in the VIPER repository. Differential expression (DE) analyses was performed with the DESeq2 R package ^68^, using supervised analysis based on gene expression levels (counts from STAR) and cutoffs of FDR-adjusted p-value (padj) < 0.05 and log2 fold-change > 1. DHT treatment of LNCaP for 4 or 16 hours was compared to vehicle treatment (two replicates each). We called loops at FDR 1%, and annotated genes to loops, following the procedure described below. The total number of loops called in 4hrs and 16 hrs were 98,960 and 183,958, respectively. The total number of genes annotated in 4hrs and 16 hrs HiChIP loop data were 17,649 and 20,899, respectively. We considered only the genes in common between LNCaP and 4hr sample (Ngenes = 15630), and between LNCaP and 16hr sample (Ngenes = 16979).

### H3k27ac HiChIP in LNCaP

HiChIP was performed mainly following the procedure described in ^61^, trypsinized 10 million LNCaP cells were fixed with 1% formaldehyde at room temperature for 10 minutes and quenched. Sample was lysed in HiChIP lysis buffer and digested with MboI (NEB) for 4 hrs. After 1hr of biotin incorporation with biotin dATP, the sample was ligated using T4 DNA ligase for 4 hrs and, chipped with H3K27ac antibody (DiAGenode, C1541019) after chromatin. Reverse crossed IP sample was pulled down with streptavidin C1 beads (life tech) treated with Transposase (illumina) and was amplified with reasonable cycle numbers based on the qPCR using 5cycle-preamplified library. Library was submitted sequenced using 150-base pair end reads on the Illumina platform (Novogene). All of these libraries were generated using the 4-bp cutter MboI restriction fragment. The raw fastq files (paired-end data) were first trimmed to remove adaptor sequences using Trim Galore ^69^. HiC-Pro version 2.9.0 ^70^ was used to align the reads to the hg19 human genome, assign reads to MboI restriction fragments, remove duplicate reads. We used the following options: MIN_MAPQ = 20, BOWTIE2_GLOBAL_OPTIONS = --very-sensitive --end-to-end --reorder, BOWTIE2_LOCAL_OPTIONS = --very-sensitive --end-to-end --reorder, GENOME_FRAGMENT = MboI_resfrag_hg19.bed, LIGATION_SITE = GATCGATC, LIGATION_SITE = 'GATCGATC', BIN_SIZE = '5000'. All other default settings were used. The HiC-Pro pipeline selects only uniquely mapped *valid* read pairs involving two different restriction fragments to build the contact maps.

We applied FitHiChIP version 5.1 ^71^ for bias-corrected peak calling and DNA loop calling. FitHiChIP models the genomic distance effect using a spline fit, normalizes for coverage differences using regression, and computes statistical significance estimates for each pair of loci. The FitHiChIP loop significance model is used to determine whether interactions are significantly stronger than the random background interaction frequency. We used 49,638 regions from H3K27Ac LNCaP union of narrow and broad peaks as anchors to call loops. We used a 5Kb resolution and considered only interactions between 5kb-3Mb. We used the peak to all for the foreground, meaning at least one anchor to be in the H3K27 peak but it doesn’t have to be both. The corresponding FitHiChiP options specification is “IntType=3”. For the global background estimation of expected counts (and contact probabilities for each genomic distance), FitHiChIP can use either peak-to-peak (stringent) or peak-to-all loops (loose) loops for learning the background and spline fitting. We specified the suggested option to merge interactions close to each other to represent a single interaction when their originating bins are closer. The corresponding FitHiChiP options specifications are “UseP2PBackgrnd=0” and “MergeInt=1” (FitHiChIP(L+M)). We used the default FitHiChIP q-value < 0.01 to identify significant loops. For comparisons across replicates, we used the results not merged MergeInt=0, as suggested by the authors. The length considered was between 5kb and 3Mb. We explored reproducibility across replicates in the following way. We have processed 5 biological replicates separately using the FitHiChIP pipeline, as well as all replicates together in a dataset called “merged” made up of the combined reads across the replicates. We used the q-value < 0.01 cutoff to define high confidence loops, and compared the level of accuracy achieved by one replicate and the merged data versus our high confidence loops (i.e., the proportion of reference loops reported by one replicate that are captured at differing number of loop calls from other replicates, or from our combined library). The final number of significant loops (q-value < 0.01) considered in these analyses are those using background 0 and merged FitHiChIP settings.

### Mapping loops to enhancers and promoters

For loop annotations, we first extended loop anchors by 5kb on either side. To identify potential gene targets, we defined promoter regions around the TSS (+/− 500 bases) for 27,063 genes using RefSeq hg19 (http://genome.ucsc.edu/cgi-bin/hgTables?hgsid=694977049_xUU5i1QkIJ50dj5miBt9wkAYuxN3&clade=mammal&org=&db=hg19&hgta_group=genes&hgta_track=knownGene&hgta_table=knownGene&hgta_regionType=genome&position=&hgta_outputType=selectedFields&hgta_outFileName=knownGene.gtf); 27 genes were removed because of ambiguous positions. The longest transcript was used to define TSS based on strand. To define enhancers, we used the subset of 49,638 regions from H3K27ac LNCaP in regular media (union of narrow and broad peaks). We then labeled the promoters and enhancer regions that overlap either right or left anchors, and considered a loop as E-E if both the anchors overlap an enhancer region; a loop as P-P if both the anchors overlap a promoter region; a loop as E-P if only one anchor overlaps a promoter and the other an enhancer region; a loop as E-O or P-O if one anchor overlaps a promoter or enhancer and the other overlaps a region without an H3K27ac or a TSS (+/− 500 bases).

### Correlation of gene expression with looping

We considered only loops involving promoters (E-P, P-P, P-O). In cases where there were duplicated loops (supporting the same two anchors but in opposite directions), we summed the PETs read counts. For each gene, we considered two measures of gene connectivity: (i) the number of loops with one anchor overlapping the promoter (the opposing anchor can overlap a promoter, an enhancer, or neither (“Other”)), (ii) the sum of PET counts across all loops overlapping the promoter. We compute a Spearman correlation between the expression across genes with both measures if connectivity. For the expression of every 13,274 genes that have looping and expression data, we averaged the FPKM value across two LNCaP RNA-Seq replicates. FPKM was converted to TPM by dividing the each FPKM value by the sum of all FPKM values of the respective sample, and multiplying by 1e6. We divided the genes into ten expression bins of equal size. We compare this to the log of the counts of loops (**Figure S3**) and the log of the counts of PETs (**Figure S4B**). Spearman correlations were computed between average TPM and log of counts of loops or PETs. To estimate the amount that each loop/PET contributes to expression, we fitted a linear regression (with or without adjusting for H3k27ac): (1) expression (TPM) ~ loops/PETs; (2) expression (TPM) ~ loops/PETs + H3k27ac. The H3k27ac level per gene is extracted by overlapping H3k27ac broad peaks scores with each gene promoter. If more than one peak overlaps the region, we sum across scores. We report the value of the coefficient in the model to represent the per-loop/per-PET contribution to expression, Spearman correlation in each version of the model, and report corresponding p-values.

### Credible sets of causal variants from Prostate Cancer GWAS

We used 147 SNPs previously reported^5^ index SNPs to define regions for fine-mapping with GWAS summary statistics from ^72^ downloaded from http://practical.icr.ac.uk/blog/?page_id=8164 (N=79,148 cases and 61,106 controls) across 20,370,946 SNPs. SNPs were compared by chromosome and position to the 1000 Genomes phase 3, for allele and rsIDs matching. After filtering by MAF>=0.001, 15,818,179 SNPs remained, with 20,155 SNPs passing genome-wide significance (p<5e-08). A ~150kb window (adjusted manually) around the 147 index SNPs, which resulted in 137 regions after merging overlapping regions on chromosomes 4, 8, and 17 was considered. In 6 of the regions (chr18.51504059.51971031.rs8093601, chr1.9945583.10662959.rs636291, chr6.170305546.170805546.rs138004030, chr7.1694537.2194537.rs527510716, chr9.123429584.124429584.rs1571801, chr9.21791998.22291998.rs17694493), a pvalue of genome-wide significance (p<5e-08) was not reached, so we do not consider these regions further. Furthermore, in the MYC region (8q24, 127.6–129.0Mb) we used 174 SNPs included in the 95% credible set from the JAM fine-mapping (Supplementary Data 3 of Matejcic et al. ^11^). Therefore, the final number of PrCa risk loci considered was 130 (129 regions which we fine-mapped (described below), and one previously mapped *MYC* region). We used PAINTOR ^73^, a Bayesian statistical method, with no functional annotations and specifying a maximum of 1 causal SNP, to fine-map 129 regions (excluding *MYC* which was previously fine-mapped). We then constructed a 95% credible set for the most likely causal variants by taking the cumulative sum of the posterior probability until a cumulative 95% posterior probability was reached. The final probable causal set contained 3,243 (3,069 from PAINTOR, 174 from JAM) SNPs across 130 PrCa risk regions. 104 PrCa loci overlapped 1,953 LNCaP loop anchors and 665 genes, connecting 2,016 fine-mapped SNPs.

### SNP-heritability enrichment for loop types

To estimate enrichment of PrCa risk heritability across functional categories, we ran stratified LDSC regression using PrCa GWAS summary statistics from ^72^. First, we created custom bed tracks using H3K27ac peaks called from ChIPseq in LNCaP. We used the custom bed tracks to annotate SNPs genome-wide indicating their overlap. We computed annotation-specific LD scores using the above annotations, using the baseline model containing 53 functional annotation^42^ and estimated functional enrichments of heritability using sLDSC ^74^.

### eQTL in prostate tissues

eQTL results on two prostate-specific datasets were used: Thibodeau eQTLs in prostate normal tissue ^40^; TCGA eQTLs in prostate tumor tissue ^41^. Thibodeau dataset contains gene expression from tumor-adjacent normal prostate tissue on 471 individuals ^40^, on autosomal and X chromosomes. TCGA dataset contains gene expression on 378 prostate cancer samples, only on autosomal chromosomes ^41^. We used MatrixEQTL ^75^ and RNA sequencing and genotype data to conduct the gene expression linear regression association for each study datasets. The cis-eQTLs were computed using a window of 3Mb from the TSS around the RefSeq genes, to match the HiChIP loop analysis. For the Thibodeau dataset, 51,232,032 SNP-eQTL pairs and 15,673 genes were tested, and a Bonferroni threshold of 0.05/51,232,032 was used to define “eQTLs” and “eGenes”. After the Bonferroni cut-off, there were 85,435 significant eQTLs and 4,747 eGenes. For the TCGA dataset, 219,256,418 SNP-eQTL pairs and 15,723 genes were tested, and a Bonferroni threshold of 0.05/219,256,418 was used to define “eQTLs” and “eGenes”. There were 86,555 significant eQTLs and 1,118 eGenes. Across both the datasets, we found 4,871 eGenes. To obtain the list of eQTL-eGene pairs, for each gene we selected the SNP with the most significant pvalue.

### Genes with somatically acquired mutations in PrCa

Genes somatically mutated in prostate cancer were extracted from three publications: an exome sequencing study looking at both localized and advanced prostate cancer ^56^, TableS4 “Known in prostate cancer and Recurrently altered in cancer” and TableS6 “cancer_pathways_mutation”); a study looking at recurrent alterations in primary (localized) prostate cancer ^57^, TableS1C “SigMutated” and TableS1D “genes in wide peak” - we only took the gene highlighted in bold - and TableS1E “Fusions”); and an analysis describing recurrent alterations in metastatic prostate cancer ^58^, TableS5 “SigMutated”). Together these sources identified 122 unique genes, of which 119 were in RefSeq and also considered in our analyses (missing MRE11A, FL1, WHSC1L1). 43 genes were within 3Mb of a credible risk SNP for PrCa.

### Gene expression data

Genes lists associated with gene expression were retrieved from the following sources:

1. 908 Genes generally expressed in prostate tissue from GTEx ^43^ (TPM > 100 TPM). We downloaded GTEx median gene-level TPM by tissue from the GTEx website (GTEx_Analysis_2017-06-05_v8_RNASeQCv1.1.9_gene_median_tpm.gct), and selected genes in RefSeq with a TPM>100.
2. 803 genes from tissue-specific expression (top 10% t.stat from ^76^). We downloaded the file from (https://data.broadinstitute.org/alkesgroup/LDSCORE/LDSC_SEG_ldscores/tstats/GTEx.tstat.tsv) which contains t-statistics comparing each gene in the particular tissue to all other tissues using 24,842 genes in GTEx. We used only the 18,026 RefSeq genes. Then, we ordered the genes based on absolute value of the t statistics in prostate, and took the top 10%.
3. 2,384 genes from differential gene expression tumor/normal ^77^. Differential gene regulation in prostate tissues was detected with GEPIA (http://gepia2.cancer-pku.cn/#degenes), which use the TCGA and GTEx projects databases to compare gene expression between tumor and normal tissues under Limma, both under and over expressed. We used the default thresholds of logFC of 1 and qvalue cut-off of 0.01.

### Over-representation of eQTL-eGene links in HiChIP loops

eGenes that overlapped HiChIP were considered: n=3,837 for Thibodeau, n=870 for TCGA. For every gene, we chose a random SNP from the same window that eQTL was computed from (i.e. 3Mb for Thibodeau and TCGA data), and computed how many times this overlaps a HiChIP anchor. We constructed 100 random control loop datasets by randomly flipping anchor1 (in 50% of the loops) or anchor2 (in the other 50%), and quantified eQTL-eGene that are supported by HiChIP loops in real data versus the random data. We find the empirical pvalue by comparing the proportion of eQTL-eGene supported by HiChIP loops in the actual versus the 100 artificial loop datasets.

### GWAS/eQTL overlap

We tested for colocalization of GWAS and eQTL using COLOC with default parameters ^78^. COLOC TCGA tested 7,001 genes, 11 genes (9 of the 130 regions) had PP4 >= 0.75. COLOC Thibodeau tested 7,148 genes, 42 genes (33 of the 130 regions) had PP4 >= 0.75. In total 46 unique genes had evidence of colocalization. 32 unique genes (3 in TCGA, 22 in Thibodeau, 7 in common across the two datasets) were also eGenes under a strict Bonferroni threshold (see **Methods** above). We used the multi-tissue TWAS for PrCa from ^79^: This includes 892 significant TWAS associations (TWAS.P < 0.05 / 109170) in 45 tissues covering 217 genes (N = 4,458), including normal and tumor prostate. Restricting to only the genes in RefSeq, this includes 651 TWAS associations in 170 genes. We restrict to prostate specific results, specifically the results in "TCGA.PRAD_SP.TUMOR", "TCGA.PRAD.TUMOR", "GTEx.Prostate", which includes 190 signals and 74 unique genes, and 42 were also eGenes under a strict Bonferroni threshold. In total 101 unique genes had evidence from COLOC or TWAS, and 74 were also eGenes. 29 genes (COLOC) and 40 (TWAS) were also HiChIP genes (n=17,690), and 24 genes (COLOC) and 26 (TWAS) were also HiChIP genes looping to a 95% credible SNP (n=665) (Table 1).

### CRISPR/dCas9-mediated repression and gene expression analysis

In order to create stable dCas9-KRAB expressing cell line LNCaP cells were infected with lenti-KRAB-dCas9-blast (Addgene, 89567) and selected with 6 µg/ml blasticidin for two weeks. sgRNAs were designed according to the "NGG" protospacer adjacent motive (PAM) restriction and gRNA efficiency score was calculated and ranked ^80^. Non-human genome targeting negative control and HPRT1 promoter targeting positive control sgRNAs were also selected. sgRNA cassettes were synthesized (Integrated DNA Technologies) and cloned into lentiGuide-Puro (Addgene, 52963) vector. All sgRNA sequences are listed in **Table S10**. LNCaP cells stably expressing KRAB-dCas9 were then subsequently infected with sgRNA vectors and selected with 2 µg/ml puromycin for five days. For gene expression, qRT-PCR 500 ng total RNA (Macherey-Nagel) was reverse transcribed (High Capacity Reverse transcription kit, LifeTechnologies) and cDNA was diluted (20x). SYBR Green assay was performed on Light Cycler 480 instrument (2x Probe Master Mix, Roche). All primer sequences are listed in **Table S11**. Relative gene expression was calculated based on the ddCT method ^81^. Each sample was measured by two biological and technical replicates. *GAPDH1* gene was used as housekeeping genes to normalize the samples.

### CRISPRi data and analysis

The ddCT method ^81^ was used to determine relative gene expression alterations from CT values of the qPCR. Briefly, for each sample, the average of the housekeeping gene (*ACTB*) was used to calculate the dCT values for each of the three technical replicates, and the average dCT values were computed for each sample. After this, using the control sample average, the ddCT values were computed for each replicate. This shows the relative deviation of each sample from the control condition. The expression values for each condition was then computed using this formula: exp = 2^-ddCT. Finally, To compute the effects and the SEs, the averages of the expression values for each sample were combined across the two biological replicates using a fixed effect meta-analysis. We then used a t-test to compare each sample to the control.

## Data Availability

We provide HiChIP interactome maps integrated with GWAS and eQTL information generated as a resource to the research community to investigate PrCa GWAS mechanisms.

## Acknowledgements

CG has reveived funding from the European Union’s Horizon 2020 research and innovation programme under the Marie Skłodowska-Curie grant agreement No 754490 – MINDED project.

## Author Contributions

CG, JHS, TS, MKF, AG, NM, BP, MLF contributed to the idea and design of the study. CG, TS, MKF, RDJ, SCB, NM performed the analyses. JHS performed the HiChIP experiment. SS performed the CRISPRi experiment. CG, JHS, TS, MKF, AG, NM, BP, MLF drafted the manuscript. All authors provided critical input on the analyses and the drafted manuscript.

