## Supplementary Figures for "H3k27ac-HiChIP in prostate cell lines identifies risk genes for prostate cancer susceptibility"

**a.**

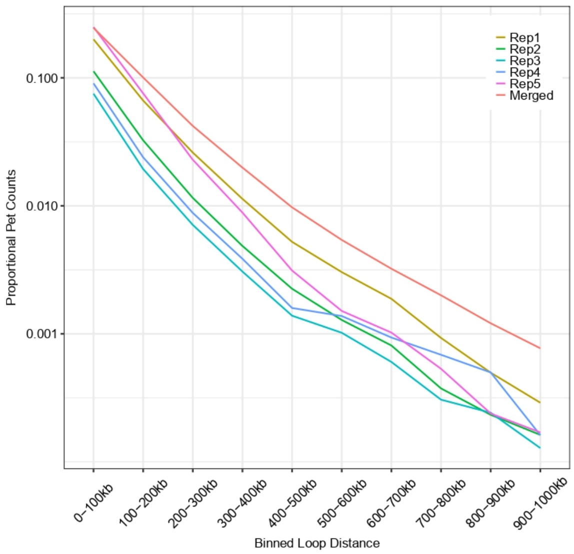


**b.**

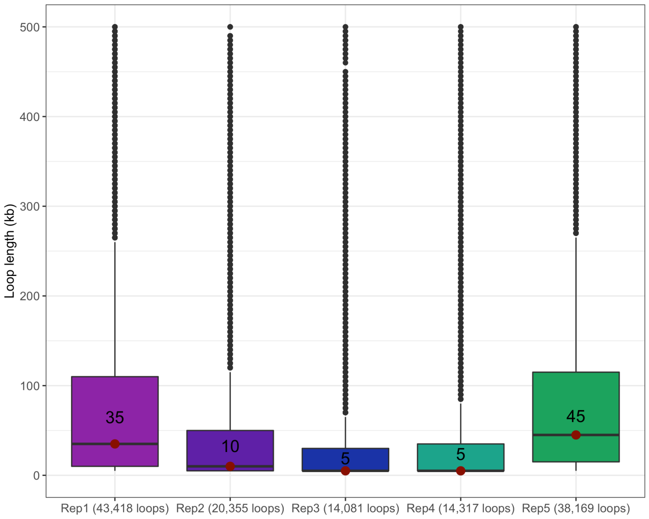


**c.**

**Supplementary Figure 1: FitHiChIP loops across 5 replicates.**

**a.** Distribution of PETs per sample (adjusted for total number of PETs in sample) as a function of the distance separating the anchors. X-axis illustrates bins of loop lengths (i.e., distance separating the anchors), y-axis, the “Proportional Pet Counts”, is the proportion of PETs divided by total number of PETs in the sample for each bin. At shorter contact distances, there are higher read counts and thus increased power.

**b.** Distribution of loop lengths across the 5 replicates at FDR 1%. The boxplots show the distribution of the loop length for each replicate. The x-axis label contains in brackets the number of loops considered (at FDR 1%) for each replicate. The box represents the interquartile range for each sample (upper and lower quartiles), the median is marked by a vertical line inside the box and written above the boxes, and the whiskers extend to the highest and lowest observations.

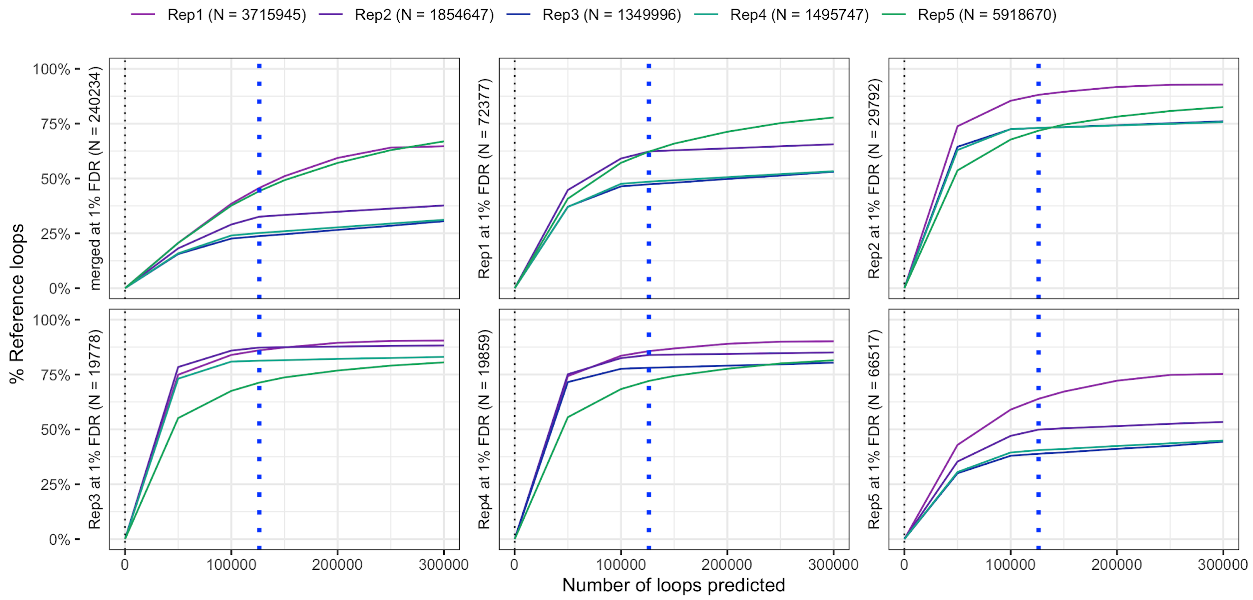


**Supplementary Figure 2: FitHiChIP reproducibility across the 5 replicates.**
For each replicate in turn, we consider the reference loops reported at FDR 1% under background 0 (no merging), and we asked what fraction of reference loops are captured by one replicate at differing number of loop calls from other replicates. The x-axis represents a differing number of loop calls from the reference data. The y-axis indicates the percent of loops of the reference data that are also called in the other replicate. The vertical dotted line highlights the n=126,280 called loops (final number of loops used in the main text). When comparing across replicates, we used the FitHiChIP setting background 0 (P2PBckgr_0) without any merging ("UseP2PBackgrnd=0" and "MergeInt=0", see **Methods**).

**a.** **b.**

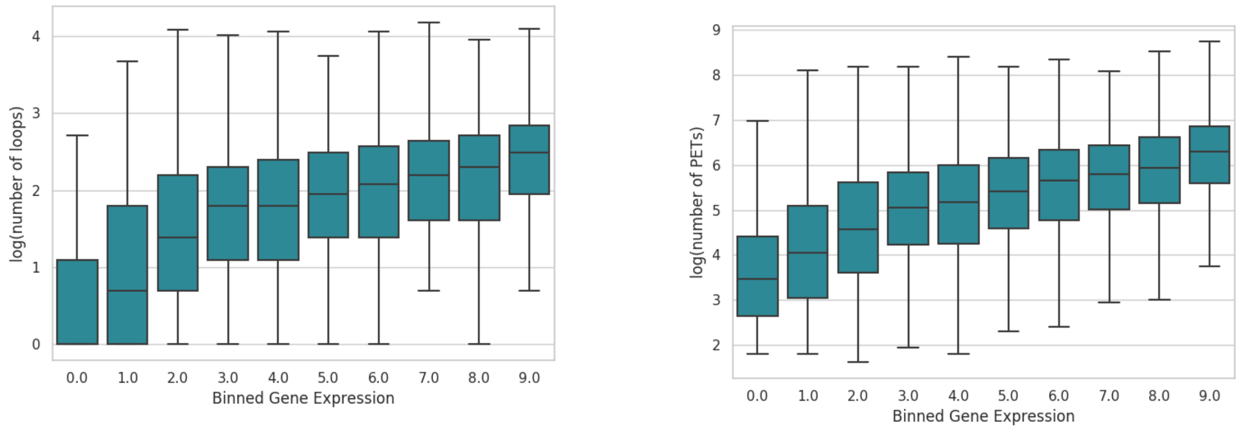


**Supplementary Figure 3. Gene connectivity and gene expression in LNCaP.** We took the union of 17,690 genes with looping counts in LNCaP and 20,114 genes with expression counts in LNCaP, dropping all genes that do not have both looping and expression information. We binned the remaining 13,274 genes into deciles (1,327 genes per decile) **a.** X-axis is the binned gene expression (FPKM) of the LNCaP genes; Y-axis represents the number of loops. **b.** X-axis is the same as the previous figure; Y-axis is the sum of PETs (b) supporting each gene in LNCaP.

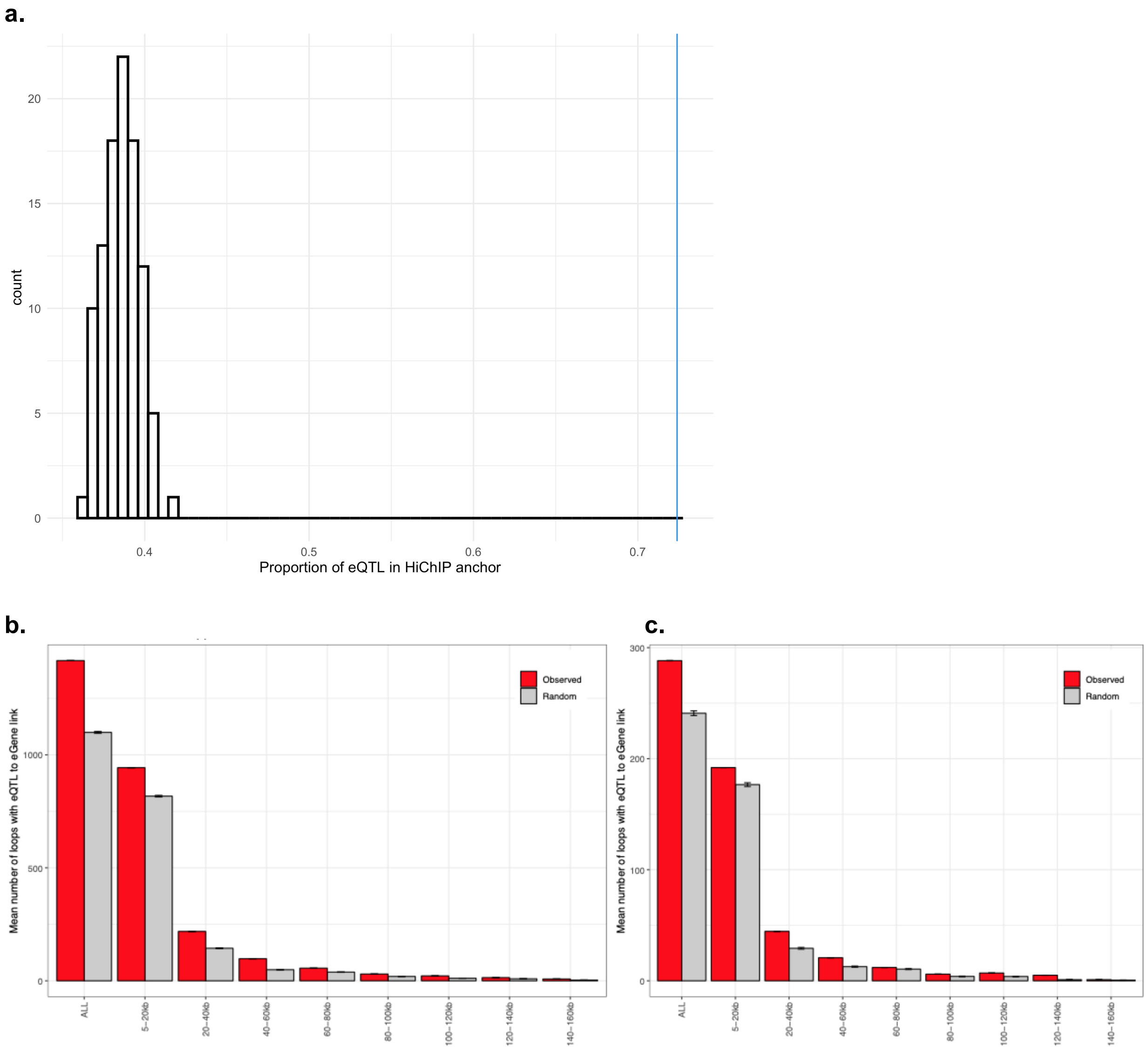


**Supplementary Figure 4: Over-representation of eQTL-eGene links in HiChIP loops. a.** Observed proportion of eQTLs falling within HiChIP anchors is compared to random. The vertical line points to the observed proportion of eQTLs falling within a HiChIP anchor. We compare this to the proportion of 100 SNPs randomly sampled within 3Mb of the promoter that are falling within a HiChIP anchor. Empirical pvalue is found by comparing the number of times the proportion from the random set is greater than the observed proportion. **b.** Number of HiChIP loops that are supported by an eQTL-eGene link in actual versus control loops across bins of loop lengths, using Thibodeau data for eQTLs. **c.** Same analysis as above using TCGA data for eQTLs. We constructed 100 control loops by randomly flipping anchor 1 or anchor 2, and identified the proportion of eQTL-eGene links falling within HiChIP anchors in the observed data compared to the random data. Using the same procedure as above, empirical pvalue is found by comparing the number of times the proportion from the random set is greater than the observed proportion.

**a**

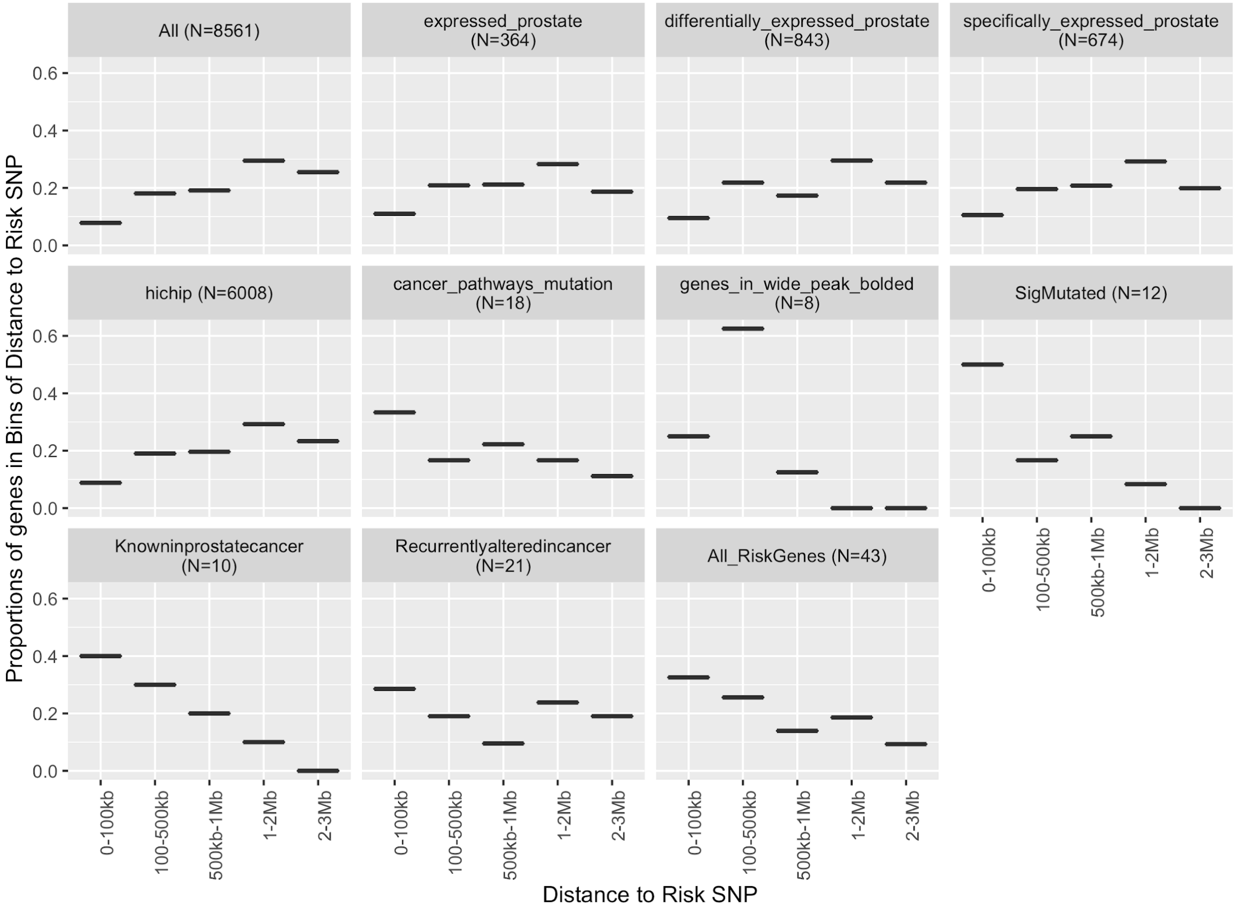


**b.**

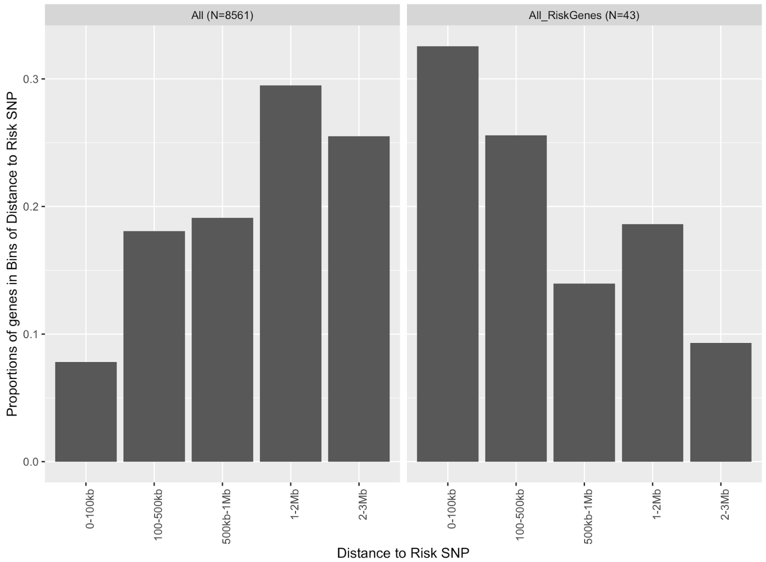


**Supplementary Figure 5: a.** Proportions of genes with respect to distance from 95% credible SNP sets, using different sets of genes. We first computed the distance between the promoter of each gene in RefSeq at a maximum distance of 3Mb (8561 genes), and 95% credible SNP set.

We then looked at proportion of genes falling in bins of the computed distances using different sets of genes (see Methods for more details on list of genes): “Expressed_prostate” (364) are the genes generally expressed in prostate tissue from GTEx (TPM > 100 TPM), “Differentially_expressed_prostate” (843) are genes differentially expressed in tumor/normal, “Specifically_expressed_prostate” (674) are the genes that have tissue-specific expression; “hichip” (6008) are the genes that are also covered in HiChIP data and have loops called at FDR 1%. We then extracted a list of risk genes previously associated with PrCa risk from three publications falling in the following five categories: cancer_patways_mutations (18), genes_in_wide_peak_bolded” (8), “SigMutated” (12), “Knowninprostatecancer” (10), “Recurrenlyalterdincancer” (21). Finally, “All_RiskGenes” (43) includes the union of the previous five categories of genes previously associated with PrCa risk. **b. Proportion of genes residing close to PrCa 95% credible risk SNP. “**All” included all genes in RefSeq considered in this paper that are within 3Mb of a PrCa credible risk SNP; “All_Risk Genes” considered genes that have been previously reported as PrCa risk genes that are within 3Mb of a PrCa credible risk SNP.
